# Increased orbitofrontal connectivity in misophonia

**DOI:** 10.1101/2020.10.29.346650

**Authors:** Leonardo Cerliani, Romke Rouw

**Affiliations:** Dept. of Psychology, Brain and Cognition, Nieuwe Achtergracht 129-B, 1018WT, University of Amsterdam; Dept. of Psychiatry, Amsterdam Medical Center, Meibergdreef 5, 1105AZ, Amsterdam, The Netherlands

## Abstract

For individuals with misophonia, specific innocuous sensory stimuli - such as hearing another person chewing or breathing - evoke strong negative emotional and physiological responses, such as extreme anger, disgust, stress and anxiety. Instead people with misophonia do not experience or display atypical reactions to generic aversive sounds such as screams or nails scratching on a blackboard. Misophonia appears to be unrelated to neurological trauma or hearing deficit, and features a characteristic developmental pattern. Its aetiology is currently unknown.

The few previous fMRI studies on misophonia showed that sufferers feature increased dorsal anterior insula activity during trigger vs. generic aversive sounds. While this effect likely reflects the saliency associated with the perception of trigger sounds in people with misophonia, in the present fMRI study we investigate the neural mechanisms underlying the emotional reaction to trigger stimuli. To this aim, we probe the task-dependent connectivity of mid-cingulate, medial premotor and ventrolateral premotor cortex. We observe that only in participants with misophonia the presentation of trigger audio-visuals prompts an increased interaction of these three brain regions with the lateral orbitofrontal cortex. This brain region is crucial for behavioural inhibition mediated by cognitive and emotional content (such as in reward-reversal learning) and is part of the temporo-amygdala-orbitofrontal network, which integrates visceral and emotional states with cognition and behaviour. We also observe that in people with misophonia trigger sounds prompt a significant increase in the interaction between mid-cingulate and the primary auditory cortex.

Our study replicates previous results and expands the network of brain regions involved in misophonia. The involvement of the orbitofrontal cortex suggests a defective functioning of high-order integrative processes allowing the reappraisal of experience-dependent negative emotional association with harmless sensory stimuli, and sheds light on the mechanisms underlying the compulsive nature of the misophonic reaction. The increased interaction, rather than the overall activity, of the primary auditory cortex with the mid-cingulate supports the hypothesis that the emotional response in misophonia is subserved by an indirect auditory-limbic pathway processing the subjective valence of specific sounds, rather than their physical properties alone.

## Introduction

Misophonia is a recently described neurobehavioural condition where specific innocuous ‘trigger’ sounds - typically human-generated sounds such as chewing, sniffing or breathing - evoke a very strong negative emotional response which manifests itself with anxiety, avoidance, deep anger and occasionally aggressive reactions (Jastreboff and Jastreboff, 2001; Edelstein *et al.*, 2013; Johnson *et al.*, 2013; Schröder *et al.*, 2013; Wu *et al.*, 2014). These emotional and behavioral patterns have detrimental effects on the social and professional life of the sufferers, potentially leading to extreme behavioural responses such as avoiding any gatherings, erupting in angry outbursts during meals, or experiencing extreme difficulty in focussing their attention on the workplace (Rouw and Erfanian, 2018; Jager *et al.*, 2020).

The emotional responses observed in misophonia occur compulsively, despite sufferers being aware of the disproportion of their emotional reaction with respect to the nature of the trigger stimuli, and regardless of the volume of trigger sounds. The latter feature clearly distinguishes misophonia from hyperacusis (Jastreboff and Jastreboff, 2001; Tyler *et al.*, 2014), qualified as a hypersensitivity to the physical properties of any sound (e.g. loudness, frequency spectrum), although a comorbidity between the two conditions is common (Jastreboff and Jastreboff, 2015). However misophonia can also not be qualified as hyperresponsivity to the subjective valence of generally aversive sounds, as sufferers have normal autonomic responses and report experiencing different feelings when hearing generic aversive sounds (e.g., baby crying, fire alarms) and trigger sounds (Kumar *et al.*, 2017). While there is evidence of comorbidity with obsessive-compulsive and mood disorders, it has been suggested that misophonia can be distinct from a secondary manifestation of a major neuropsychiatric disorder (Jager *et al.*, 2020). Currently misophonia is however not yet contemplated by psychiatric diagnostic manuals, and its exact definition is still debated (Kumar and Griffiths, 2017, Schröder *et al.*, 2017*b*).

The originally proposed neurological model for misophonia suggests an atypically enhanced interaction between sensory - especially auditory - and limbic systems (Jastreboff and Hazell, 2004; Edelstein *et al.*, 2013; Jastreboff and Jastreboff, 2015). Recently, neuroimaging studies began to identify the network of brain regions which interact to generate the emotional and behavioural response to trigger stimuli in misophonia. These studies highlighted the presence of an increased activity in the anterior dorsal insula during trigger stimuli compared to generic aversive sounds (Kumar *et al.*, 2017). While the insula represents an important node for the low-level integration of sensory and emotional, as well as autonomic and cognitive information (Mesulam and Mufson, 1985; Gogolla, 2017), increases in its activity - and especially in its anterior division - can be elicited by any event impacting the emotional and autonomic homeostasis of the organisms, such as pain, social exclusion, gambling, (Critchley, 2005; Seeley *et al.*, 2007; Craig, 2009; Kurth *et al.*, 2010; Nieuwenhuys, 2012). Thus the increased insular activity recorded during trigger stimuli cannot explain *per se* why sufferers of misophonia react to these sounds differently from aversive sounds or other emotionally-loaded stimuli (Craig, 2010; Cauda *et al.*, 2012; Perini *et al.*, 2018).

Importantly in the same study (Kumar *et al.*, 2017) it was shown that the presentation of misophonic stimuli triggered an increased interaction (functional connectivity) between the anterior dorsal insula and the default-mode network (Fox *et al.*, 2005). Instead no alteration was found in the activity or connectivity of the auditory cortex, and only a marginally significant effect was found in the functional connectivity of the amygdala (Kumar *et al.*, 2017). This important evidence suggests that in misophonia the extreme emotional reaction to trigger stimuli is not subserved by direct sensory-limbic circuits responding to the physical properties of the sound alone, but rather mediated by high-level cognitive associations between the stimulus and its subjective valence.

Given that the dorsal anterior insula and the default mode network are respectively involved in estimating the subjective valence of the sensory stimuli (Craig, 2009), and in internally directed cognitive processes - such as social cognition and autobiographical memory (Margulies *et al.*, 2016) - these findings shed light on the perceptual processes involved in misophonia, but they leave open the question of which brain regions and neural networks are involved in the generation of the emotional response to trigger stimuli.

Besides the anterior insula, other brain regions show increased activity during trigger stimuli in a whole-brain corrected analysis (Kumar *et al.*, 2017), namely the mid-cingulate cortex (MCC) - which together with the anterior insula defines the saliency network (Seeley *et al.*, 2007) - the supplementary motor area (SMA) and the ventrolateral premotor cortex (vlPMC). All these regions are likely to play an important role in the misophonic response, by prompting an autonomic-driven emotional and behavioural response such as the urge to avoid or react to the trigger sounds (Schröder *et al.*, 2013; Perini *et al.*, 2020). However, the activity of these brain regions in misophonia has not yet been further investigated.

In the present study we use fMRI to systematically investigate the presence of alterations in the task-dependent connectivity across all brain regions whose activity is enhanced in misophonia when experiencing trigger stimuli. To this aim, we contrast the brain response to misophonic triggers with that associated with sounds generally perceived to be aversive (e.g., nails on chalkboard) but not eliciting atypical emotional response in people with misophonia. This contributes to define the network of regions specifically involved in this complex association of innocuous sensory stimuli with extremely negative emotional responses in misophonia. We expected this investigation to bring further evidence to the hypothesis that misophonia is grounded on high-order cognitive associations, rather than in an atypical perception of the trigger sounds. Another non-secondary aim of the present study was to replicate previous fMRI results, given the current paucity of neuroimaging studies on misophonia.

## Methods

### Participants

In a preliminary behavioural study, we examined a large number of individuals with misophonic complaints (N=232), measuring severity and nature of the misophonic complaints (Rouw and Erfanian, 2018). In this relatively young research field, an agreed upon diagnostic tool for misophonia is still lacking. A number of questionnaires have been proposed by researchers and clinicians to measure the nature and severity of misophonic complaints (Fitzmaurice, 2010; Edelstein *et al.*, 2013; Schröder *et al.*, 2013; Wu *et al.*, 2014; Bauman, 2015; Dozier, 2015). The commonalities across these questionnaires highlights which characteristics are currently viewed as key to the condition of misophonia. The misophonia symptoms severity (MSS) questionnaire we used for the selection of participants is a collection of 15 items measuring at an interval scale the common features of misophonia identified in previously proposed questionnaires (see Supplementary materials).

Participants rated statements, such as *I feel angry when I hear/see a trigger* on a 5-point scale, ranging from “Strongly disagree” (1) to “Strongly agree” (5). A misophonia symptoms severity (MSS) score was calculated as the mean of all 15 items. This preliminary study showed that our questionnaire has high scale reliability (Cronbach's alpha 15 items; α = 0.97; Guttman’s lambda-2 α = 0.97), and created a strong basis for selecting participants to invite for the neuroimaging experiment. For the neuroimaging study, we invited participants with very high MSS scores (not lower than 3 in any item), and did not meet the following exclusion criteria: visual impairments, age below 18 or above 60 years old, presence of non-removable metal objects in the body, (possible) pregnancy and claustrophobia.

We checked the presence of misophonia complaints in control subjects with an interview as well as with the MSS questionnaire. In total, thirty-nine participants were included in the present analyses; 19 individuals with misophonia (15 female and 4 male, mean age 31 years, range 18-58) and 20 controls (15 female and 5 male, mean age 29 years, range 19-53). Groups were matched by gender and age (T_(37)_ = 0.4, p = 0.69). The mean score on the MSS questionnaire in the misophonia subject group was 3.9 (score range: 3.2 to 4.5). The mean score for the control group was 1.45 (scores range: 1.0 to 2.2). The MSS scores were significantly higher in people with misophonia (T_(37)_ = 18, p < 0.0001).

### fMRI paradigm and behavioural ratings

We used audio-visual stimuli, rather than only sounds as in (Kumar *et al.*, 2017), to better approximate real-life conditions (Schröder *et al.*, 2019). Misophonic stimuli (hereafter: “trigger stimuli”) were 12-second audio-visuals in which an actor produced sounds that were known to elicit a particularly negative emotional response (anger, disgust) in our participants with misophonia, such as chewing an apple, heavily inhaling with the nose and clearing the throat. These trigger stimuli were contrasted with clips presenting generically aversive sounds (e.g. nails on a chalkboard) and with clips in which the actor created neutral sounds (e.g. peeling an apple, shaking a bottle of water).

For each of three categories of audio-visuals - trigger, aversive, neutral - we presented inside the scanner six clips of 12 sec. each. Each clip was preceded by a short text description of 2 sec. and followed by a rating time of 6 sec. Audio-visual presentation and rating time were separated by blank screens of variable durations from 1 to 5 seconds.

After the presentation of each audio-visual, participants were asked to evaluate their own emotional reaction (positive or negative) to that particular audio-visual clip. They responded by moving a cursor on the screen (by pressing the left or right button on the response box) along a dotted line, representing an 11 points Likert scale ranging from minus 5 (very negative) to 0 (neither negative nor positive) to plus 5 (very positive). Since in the neuroimaging analyses a more negative rating is a proxy for increasing unpleasantness associated with different stimuli, in the results reported below the ratings are reversed, so that higher scores reflect more negative feelings and increasing unpleasantness (see Figure 1). We choose to use a rating scheme that would also cover positive feelings to avoid biasing the ratings towards negative feelings also for stimuli which might have been perceived as neutral or positive.

**Fig. 1.**
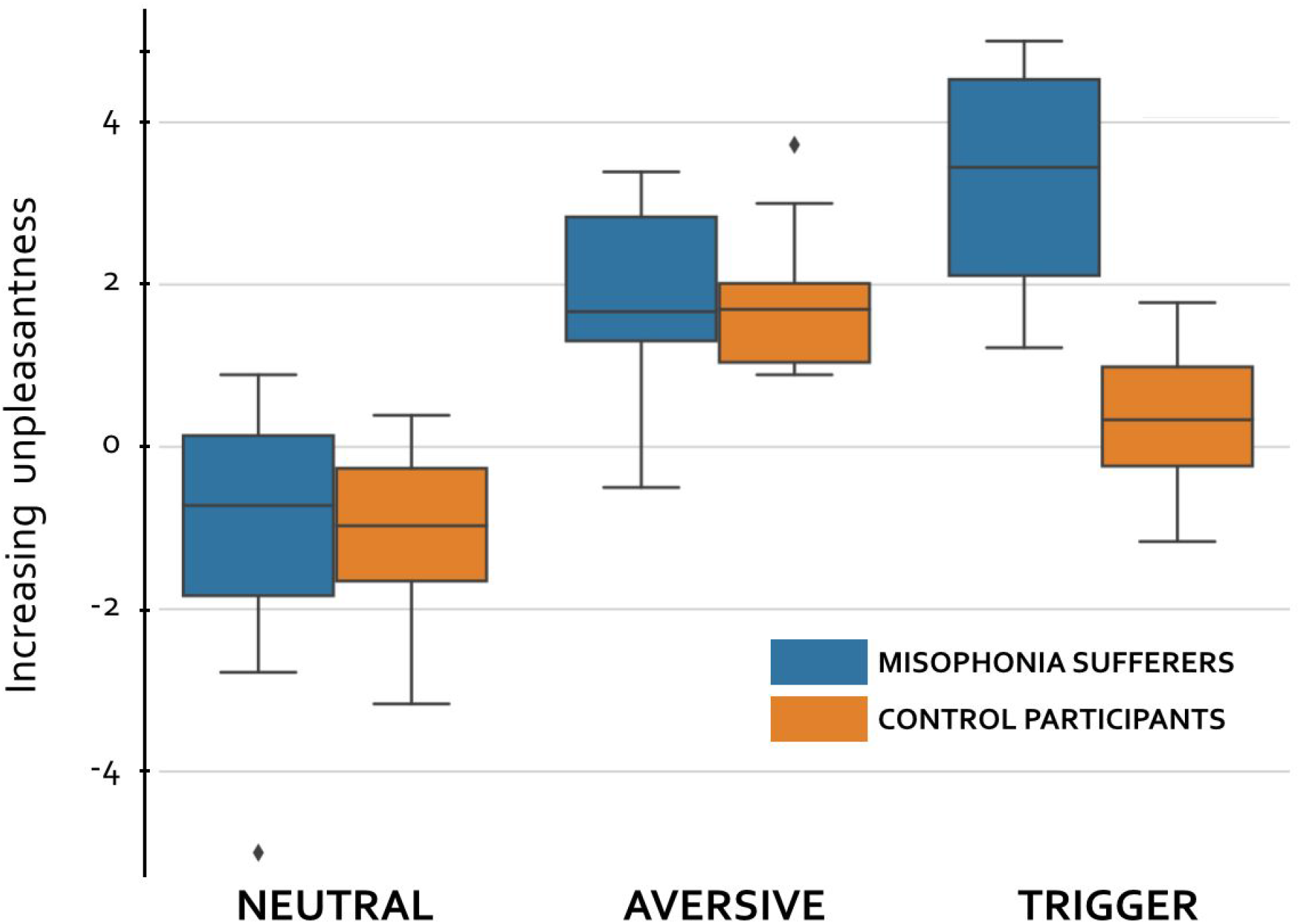
Audio-visual ratings during the fMRI task. Higher values indicate that the stimuli are perceived as more unpleasant. For each participant, we considered the average rating across stimuli in each category (Neutral, Aversive, Trigger). Boxplots display the median and the first and third interquartile. Diamonds indicate participants whose rating for that category was bigger than 1.5 times the interquartile range, the latter represented by the whiskers.

Each participant completed a sequence of 3 runs for the entire paradigm. The order of the clips was different across the three runs, but identical for all subjects within each run. Blank screen time always amounted to 108 sec. per run, but the order of the breaks was randomized for each participant and each run. In total, each run lasted 468 sec.

### MRI data acquisition

MRI images were acquired at the Spinoza Center for Neuroimaging (Amsterdam, NL) on a Philips 3T Achieva scanner equipped with a 32-channel head coil. For each participant, fMRI images were acquired with a Philips T2* FEEPI sequence with the following parameters: repetition time (TR) = 2.0 sec, echo time (TE) = 27.63 ms, flip angle = 76.10°, voxel size = 3×3×3 mm, slice gap = 0.3 mm, 37 axial slices, FOV=240×121.8×240. For each of the three fMRI runs, 234 whole-brain images were acquired. Participants were instructed to stay still and relax their head. Head movements were restrained with padding cushions around the head within the head coil. In the same session, an anatomical T1w image covering the whole brain was acquired, with the following parameters: 220 axial slices, voxel resolution = 1 mm^3^ isotropic, TE = 3.86 ms, TR = 8365 ms, flip angle = 8°.

### Analysis of movie ratings

The analysis of movie ratings was carried out in JASP (JASP Team, 2020). Besides including these ratings in our fMRI model, we examined this data alone to test for the expected difference in ratings between groups only for trigger stimuli. For each participant, ratings were averaged across audio-visuals of the same category, to obtain one mean rating for trigger, aversive and neutral movies per participant.

### fMRI data analysis

#### Preprocessing

Analysis of fMRI and T1w data were carried out using the [FSL FMRIB Software Library](https://fsl.fmrib.ox.ac.uk/fsl/fslwiki/) (Woolrich *et al.*, 2009; Jenkinson *et al.*, 2012). The T1w image of each participant was skull-stripped using bet2 (Smith, 2002) and the results visually inspected. The fMRI data of each participant was preprocessed with the following parameters: motion correction using MCFLIRT (Jenkinson *et al.*, 2002); slice-timing correction; non-brain removal (Smith, 2002); spatial smoothing with a Gaussian kernel of FWHM 5 mm; grand-mean intensity normalisation of the entire 4D dataset by a single multiplicative factor; high pass temporal filtering (Gaussian-weighted least-squares straight line fitting, with sigma=50.0s).

#### Standard group-by-task fMRI analysis

The effect of each task of interest - trigger, aversive or neutral audio-visuals - on brain activity was modelled as a boxcar spanning the duration of the audio-visual, convolved with a standard haemodynamic response function. The amplitude of the boxcar reflected the (mean corrected) increasing perceived unpleasantness - derived from the behavioural ratings - that the participant associated with each audio-visual. Therefore this predictor modelled the effect of the audio-visual, modulated by the associated unpleasantness rating (Poline *et al.*, 2007). We also generated confound predictors for the rating period, button press during the rating period (orthogonalized with respect to the rating period predictor), text presented before the audio-visuals. The temporal derivatives of these 6 predictors were also added to the model, as well as 18 additional confound predictors modelling motion parameters.

We then defined contrasts of parameter estimates (COPEs) to compare brain activity between each category of stimuli: Trigger > Aversive; Trigger > Neutral; Aversive > Neutral. For each participant, the contrast of parameter estimates maps (COPEs) were averaged across the three runs in a 2nd-level analysis using a fixed effect model. Group-level effects were then estimated in a 3rd-level mixed-effect analysis using FLAME-1 (Woolrich *et al.*, 2004) including the average COPEs from the 2nd-level analysis of each participant. Z image statistics were thresholded using clusters determined by Z > 3.1 and a corrected cluster significance (using Gaussian Random Fields) of p=0.05 (Worsley, 2001).

#### Psychophysiological interaction

Psychophysiological interactions (PPI) are task-dependent variations in the functional connectivity - that is, in the temporal synchronization - between the activity of two brain regions (Friston *et al.*, 1997).

The standard group-by-task fMRI analysis detailed above detects brain regions where an *average increase in activity* - across all the time of the audio-visual - is observed when contrasting the task of interest (trigger audio-visuals) with a control condition (aversive or neutral audio-visuals). Instead the PPI analysis probes the *synchronization of brain activity between two brain regions due to the task*. This allows to differentiate regions which interact with each other during the task, from regions which, although both activated by the task, work independently (O’Reilly *et al.*, 2012).

The results of PPI can be interpreted as a modulatory effect of the task on the interaction between two brain regions, or as the modulatory activity of one brain region on another brain region when the latter is (or both are) processing a stimulus (See Fig. 5 in (Friston *et al.*, 1997)). Importantly, regions associated with this modulatory activity by the standard fMRI analysis, as the increase in *functional connectivity* between two brain regions is not necessarily accompanied by an increase in *brain activity* (or more precisely in the intensity of the blood-oxygen-level dependent signal, which is measured by the fMRI acquisition) during the task of interest with respect to the control condition.

PPI is practically implemented by adding two predictors to the model used in the standard analysis: (1) a physiological predictor, represented by the time course of the (seed) region of interest - usually identified in the initial standard analysis; (2) a psychophysiological predictor, represented by the element-by-element multiplication between the task predictor (psychological predictor) and the physiological predictor (Friston *et al.*, 1997; O’Reilly *et al.*, 2012; Di *et al.*, 2020).

In our study, we seeded one PPI analysis in each of the clusters where a significant increase in brain activity was detected when contrasting Trigger > Aversive stimuli in the initial standard analysis. For each cluster, the time course of the signal was extracted using a sphere of 5 mm radius centered on the local maxima, and demeaned across time points. This generated a COPE map quantifying the effect of the PPI predictor for each task (trigger, aversive, neutral). These COPE maps were then processed at the group level as in the standard analysis, to yield corrected thresholded Z image statistics of the group differences for each task. Therefore, also in this case Z image statistics were thresholded using clusters determined by Z > 3.1 and a corrected cluster significance (using Gaussian Random Fields) of p=0.05 (Worsley, 2001).

## Results

### Perceived unpleasantness in audio-visuals

Figure 1 shows the distribution of the average audio-visual ratings, across stimuli of the same category, in people with misophonia and in control participants. A two-way ANOVA with factors for stimuli and groups shows a significant interaction between the factors (F_(2,111)_ = 20.94, p < .001 ω^2^ = 0.17) as well as a main effect of both stimuli (F_(2,111)_ = 103.6, p < .001 ω^2^ = 0.47) and of group (F_(1,111)_ = 26.51, p < .001 ω^2^ = 0.074). Crucially, the two groups significantly differ only in the ratings for Trigger stimuli (T_(76)_ = 8.23, p < 2.9e-9) as misophonia sufferers rate trigger sounds as more unpleasant, while there is no significant difference in perceived unpleasantness for Aversive (T_(76)_ = 0.48, p = 0.63) or Neutral (T_(76)_ = 0.21, p = 0.83) stimuli.

### Standard task-by-group fMRI analysis

In the Trigger > Aversive contrast, people with misophonia featured significantly increased brain activity in the bilateral supplementary motor area (SMA) and mid-cingulate cortex (MCC), in early visual regions (V1/V2) and in the right ventrolateral premotor cortex (vlPMC), with the latter cluster extending in the dorsal anterior insula and adjacent frontal operculum (aIC/FOP) (Figure 2). No significantly decreased activity was observed for this contrast. These differences were not found for the Trigger > Neutral contrast at this threshold, although similar midline differences were observed with an initial cluster-forming threshold of Z > 2.3. However, we didn’t further consider the latter results as the use of this lenient threshold has recently been discouraged, since it was shown to potentially yield excessive false positives and localization issues (Woo *et al.*, 2014).

**Figure 2.**
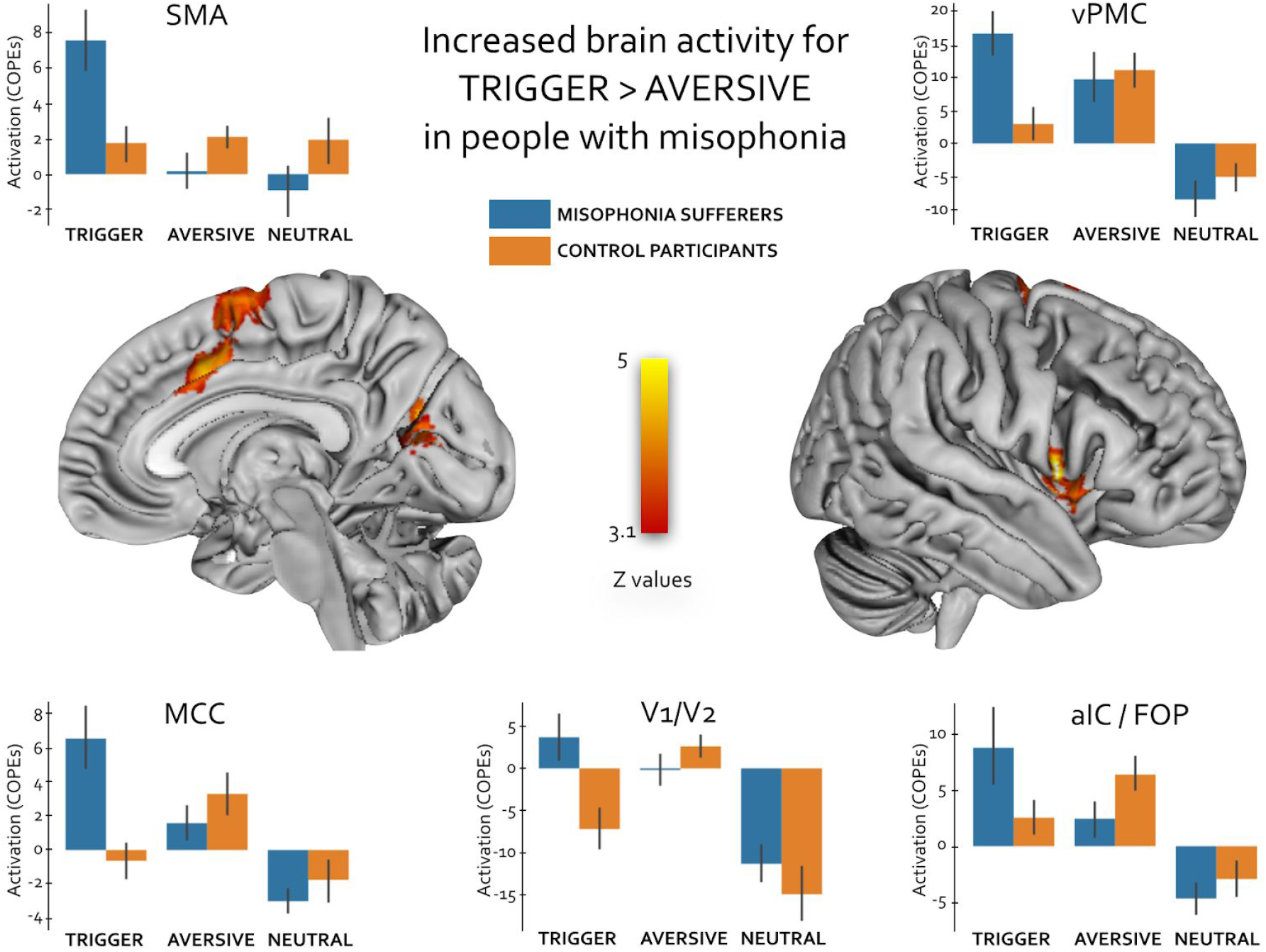
Between-group differences in fMRI brain activity for the contrast Trigger > Aversive. These images display the significant results in the contrast Misophonic participant [Trigger > Aversive] > Controls [Trigger > Aversive]. Results are whole-brain corrected with a cluster forming threshold of Z > 3.1 and corrected cluster significance of p = 0.05. Barplots display for each location the mean (+/− standard error) parameter estimates of each contrast (COPEs) in each group for all the voxels in a sphere of 5 mm radius centered on the local maxima.

### Psychophysiological interaction (PPI)

The PPI analysis showed that in people with misophonia, the presentation of Trigger audio-visuals increased not only the overall activity, but also the functional connectivity of MCC, SMA and vlPMC with several brain regions, including the lateral and medial premotor and prefrontal cortices, the primary auditory cortex (A1), and especially the lateral orbitofrontal cortex (OFC) (Figure 3). The PPI analysis for V1/V2 did not yield any significant results. The same PPI analysis for generic aversive and neutral stimuli did not yield any significant result.

**Figure 3.**
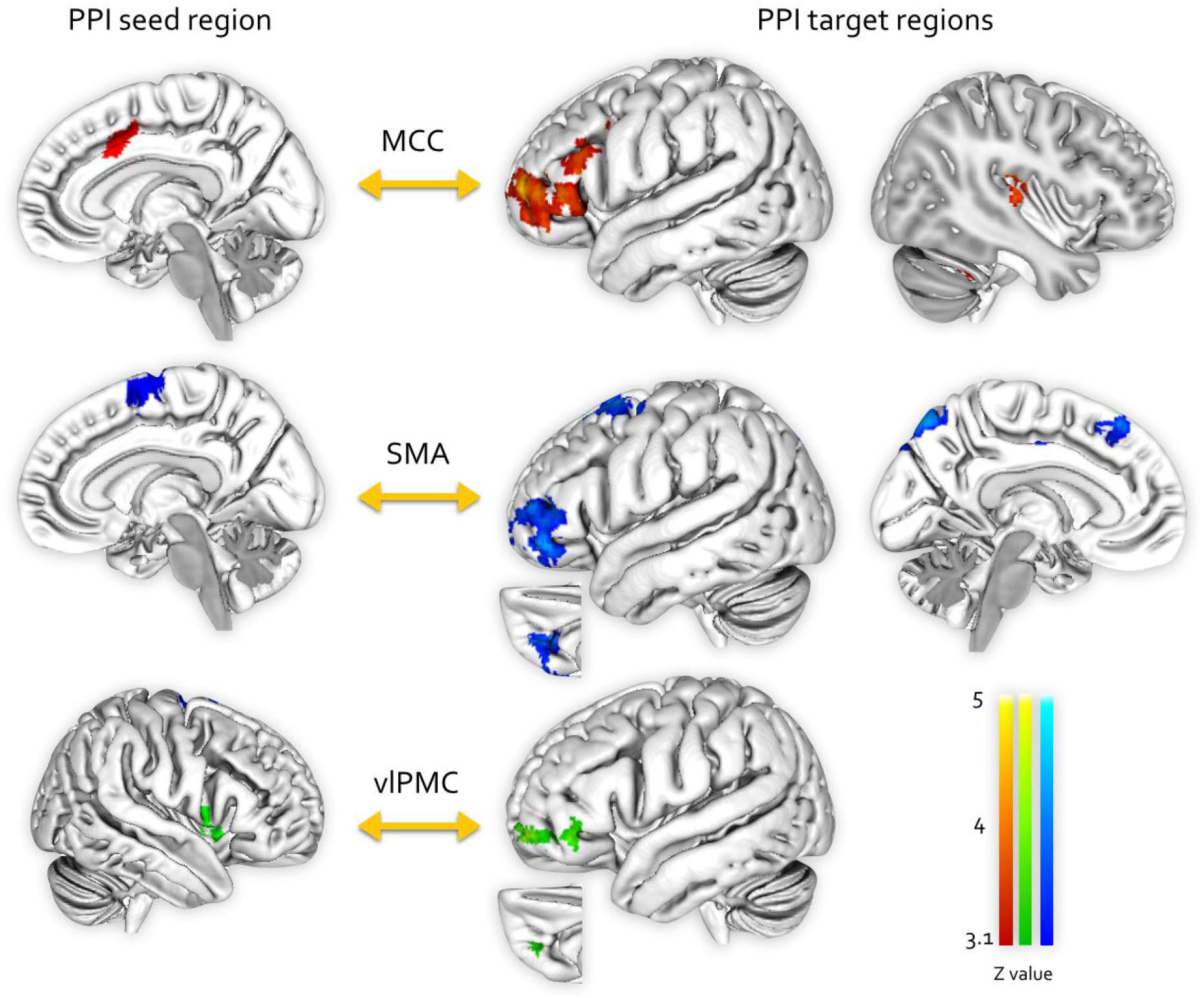
Summary of the PPI findings for trigger stimuli in people with misophonia. The presentation of Trigger stimuli prompts an increase in task-dependent functional connectivity between MCC, SMA, vlPMC and other brain regions. The seed region of the PPI analyses and the location of their significant effects in the rest of the brain (PPI target regions) are matched by color. Results are whole-brain corrected with a cluster forming threshold of Z > 3.1 and corrected cluster significance of p = 0.05. No significant PPI results were found for V1/V2. This effect was significant only for people with misophonia and only during Trigger audio-visuals. Generic aversive and neutral audio-visuals did not elicit any significant PPI effects.

Figure 4 highlights the increased connectivity between MCC, SMA, vlPMC and the lateral OFC. It is remarkable that despite the different location of the seeds, the strongest increase of their functional connectivity with any other part of the brain was co-localized in a brain region including the ventral part of the inferior frontal gyrus, the adjacent frontal pole and the basal orbitofrontal cortex. The latter regions recapitulate the frontal projections of the uncinate fasciculus (see Figure 4, bottom-right). This effect was observed only in people with misophonia and only during the presentation of trigger stimuli.

**Figure 4.**
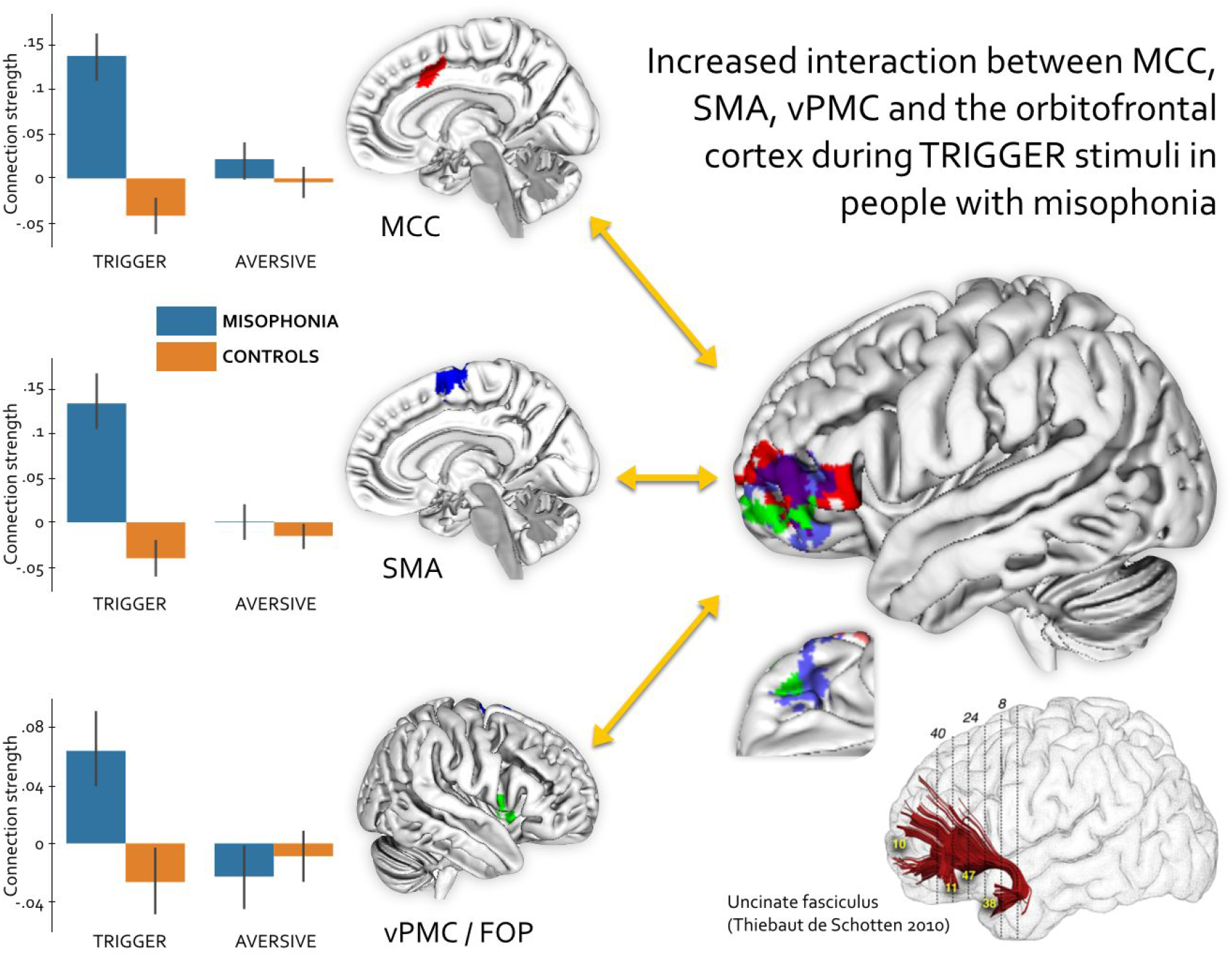
Increased task-dependent synchronization between MCC, SMA, vlPFC and the lateral OFC during Trigger audio-visuals in people with misophonia. The overlay on the brain surface in the right part of the figure highlights the significant results from the PPI analysis (Figure 3) which were located in the OFC. This cortical territory receives the fronto-lateral projections of the uncinate fasciculus (bottom-right). The seed region of the PPI analyses and the location in the brain of their significant effects are matched by color. A certain degree of transparency is applied to the result for the SMA (blue overlay) to show the overlap with the results from MCC (red) and vlPMC (green). The barplots on the left display, for each seed region, the mean (+/− standard error) parameter estimates of PPI connection strength across all the corresponding OFC voxels featuring a significant effect.

Besides the lateral OFC, another particularly interesting result in the context of misophonia is the increased task-dependent interaction between MCC and a region localized in the Heschl gyrus (primary auditory cortex - A1) and extending also to the posterior insular cortex. This effect, and the corresponding PPI parameter estimates of connection strength, are shown in Figure 5.

**Figure 5.**
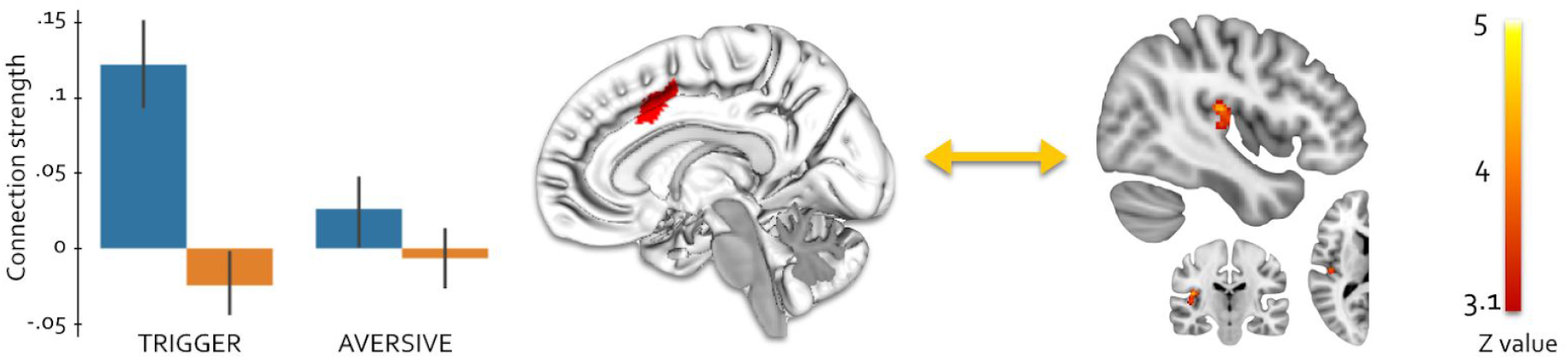
Increased interaction between MCC and primary auditory cortex (A1) - The MCC seed region is the same shown in Figure 3 and 4. The barplot on the left displays the mean (+/− standard error) parameter estimates of connection strength across all the corresponding A1 voxels featuring a significant effect.

## Discussion

Misophonia is a recently described behavioural condition (Jastreboff and Jastreboff, 2001), and so far its brain mechanisms have been investigated using neuroimaging only in three studies (Kumar *et al.*, 2017; Eijsker *et al.*, 2019; Schröder *et al.*, 2019). While the anterior insular cortex has emerged as a region of interest for the perception of trigger stimuli, much less is known with respect to the interaction among all brain regions involved in the disproportionate negative emotional reaction to the trigger stimuli which characterizes misophonia.

In the present study we first replicate previous findings describing where in the brain trigger stimuli elicit an increase in brain activity in misophonia. Then by investigating their task-dependent connectivity we report the previously undescribed involvement of the lateral orbitofrontal cortex (OFC) and of the primary auditory cortex (A1). In the following we discuss our results in comparison with previous neuroimaging evidence, and elaborate on why the atypical functional connectivity between premotor, cingulate and OFC cortices represents a good candidate to reflect the emotional reaction to trigger stimuli in misophonia.

### Comparison with previous neuroimaging studies

The first neuroimaging study on misophonia (Kumar *et al.*, 2017) highlighted a prominent increase of brain activity in the bilateral anterior dorsal insular cortex when people with misophonia were listening to trigger vs. aversive or neutral sounds. We replicated the effect of trigger stimuli on insular activity in misophonia, although not to the extent reported by (Kumar *et al.*, 2017), and only in the right hemisphere. Importantly, in our paradigm the stimuli were preceded by a short textual description of the forthcoming stimulus, to isolate the effect of the sensory content of the stimulus with respect to its verbal associations. The contrast [text description] > [audiovisual stimuli] did not result in significant group differences, however in both groups separately this contrast showed a significant higher activity in the left anterior insula (See Supplementary Figure 1). Given the predominant left hemispheric lateralization of language (Mazoyer *et al.*, 2014) as well as the left hemispheric lateralization of the insular cortex for linguistic, rather than emotional aspects of speech production (Ackermann and Riecker, 2004, 2010), our result suggest that left and right anterior insulae processed different aspects of the audiovisual stimuli - linguistic associations and emotional content, respectively - and that the misophonic reaction was selectively determined by non-verbal emotional associations. However, our paradigm was not designed to test the impact of different modalities in the generation of the misophonic response, therefore further investigation is needed to support this hypothesis.

Increased activity in the ventral anterior insula had also been previously reported when this region was manually selected *a priori*, however this result was not significant in the whole-brain corrected analysis (Schröder *et al.*, 2019). In our analysis we could not find an evidence of increased ventral insular activity or functional connectivity during trigger stimuli, even *without* appropriate correction for multiple comparison. It is important to mention that the ventral anterior insula has very different functional, connectional and cytoarchitectonic properties with respect to the dorsal anterior insula (Mesulam and Mufson, 1982; Kurth *et al.*, 2010; Cerliani *et al.*, 2012), and represents an important direct gateway of sensory and autonomic information to the limbic system (Mesulam *et al.*, 1985; Stefanacci and Amaral, 2002; Heimer and Van Hoesen, 2006; Nieuwenhuys, 2012). Instead the dorsal anterior insula represents a major node of the saliency network, together with the mid-cingulate cortex, and is involved in higher-order representations of sensory, cognitive and interoceptive information, to the extent that they are related to changes in the emotional homeostasis of the organism and to emotional awareness (Craig, 2009, 2010). The limited effect of triggers on the ventral vs. dorsal insular activity is the first element suggesting that the emotional reaction in misophonia is mediated by high-order constructs, rather than by a direct auditory-limbic connection relaying only the physical properties of the sound (volume and frequency spectrum). Further support to this hypothesis is provided - as we will discuss in the next paragraphs - by the involvement of the OFC and by a comparison between misophonia and hyperacusis.

Besides the anterior dorsal insula, contrasting trigger with aversive stimuli also resulted in significantly higher brain activity in the medial premotor cortex (supplementary motor area - SMA), mid-cingulate (MCC) and ventrolateral premotor cortex (vlPMC). The locations of the increased activity correspond to those reported by Kumar and colleagues (see Table S1 in (Kumar *et al.*, 2017)). All of these regions are primarily responsible for planning and preparing motor behaviour and are likely to be related to the urge to avoid or react to trigger sounds in people with misophonia (Schröder *et al.*, 2013). Specifically, an increased activity in the premotor cortices reflects planning or inhibiting a motor response, while the mid-cingulate cortex signals that the behavioural response is associated with an autonomic response. Related to this, we recently showed that the response associated with a basic (negative) stimulus - heat-induced pain - results in an increased interaction between the anterior insula and the mid-cingulate - proportional to the urge to move - in neurotypical participants, while this mid-cingulate connectivity is not increased in carriers of the R221W mutation (of the nerve growth factor gene), in whom a painful stimulation does not elicit an autonomic response (Perini *et al.*, 2020). Therefore the increased activity in SMA, MCC and vlPMC is arguably reflecting the neural activity involved in the emotional and behavioural reaction to trigger stimuli, rather than in their perception, as an effect of the autonomic change induced by trigger stimuli, likely associated with the increased insular activity.

One notable difference in our results with respect to previous reports is the increased activity in primary visual regions (V1/V2). This is likely due to the fact that our stimuli were audio-visuals rather than only sounds as in (Kumar *et al.*, 2017). This choice was motivated by the purpose of providing in the MRI scanner a multimodal experience of trigger sounds, similar to real-life contexts. The cause of this increased primary visual activity is however unclear. A previous study reports that autonomic changes in people with misophonia were elicited by sounds but not by corresponding images (Edelstein *et al.*, 2013), while no study yet examined whether the multi-modal aspect of trigger stimuli would contribute to amplify the emotional response in misophonia (Brout *et al.*, 2018). If the interaction between different sensory modalities plays a role in eliciting the misophonic response, audio-visual coupling could potentially be manipulated - as in the McGurk effect (McGurk and MacDonald, 1976) - to alleviate the symptoms of misophonia (Samermit *et al.*, 2019).

### Increased orbitofrontal connectivity

The most important result of our study is represented by the increased interaction between premotor and cingulate cortices with the orbitofrontal cortex, which was significant only in people with misophonia and only during trigger stimuli. Furthermore, the presentation of misophonic stimuli *per se* did not increase the overall brain activity in OFC, suggesting that this brain area is particularly involved in shaping a functional unity (synchronization) of brain areas during the misophonic response.

The lateral OFC is part of a major cortico-subcortical limbic network which anatomically connects the amygdala and the paralimbic temporal pole with the anterior insula, the cingulate cortex, the lateral OFC and the adjacent frontal pole via the uncinate fasciculus. This temporo-amygdala-orbitofrontal (TAO) network is responsible for the integration of visceral and emotional states with cognition and behaviour (Mesulam, 2000; Catani *et al.*, 2013).

Within the TAO network, the lateral OFC is involved in establishing high-level association between sensory stimuli and behavioural responses, and in particular in updating the latter according to mutated environmental conditions. More specifically, the lateral OFC is crucial in estimating the emotional consequences of a learned behavioural response to specific stimuli (Vogt, 2019*b*): lesion studies have shown that damages to this brain region impair the inhibition or correction of the behavioural response when a previously rewarded stimulus-behaviour association is not anymore rewarded (reward-reversal learning) (Rolls, 2019). This suggests that the lateral OFC activity is crucial to prevent the establishment of compulsive behavioural patterns which are not (anymore) favourable. In this respect a recent fMRI study on stop-signal inhibition on a large sample (N=1709) of healthy adolescents found that the activity of OFC - as well as of mid cingulate and dorsolateral prefrontal cortex - significantly increased in successful vs. failed behavioural inhibition (Deng *et al.*, 2017). Furthermore, a symptom provocation study in patients with obsessive-compulsive disorder (OCD) found that the lateral OFC selectively responds to OCD-relevant stimuli in comparison to generic aversive stimuli (Simon *et al.*, 2010).

In the specific context of misophonia, the increased trigger-specific functional connectivity of the lateral OFC could reflect the attempt to downregulate the emotional response which is automatically prompted by the trigger stimuli, given the role of this brain region in the reappraisal of stimuli associated with negative emotions (Banks *et al.*, 2007). The automatic response prompted by trigger stimuli might originate from previously learned association mediated by the lateral OFC, and facilitated by the hyperconnectivity between the anterior dorsal insula and the default mode network, as previously reported (Kumar *et al.*, 2017). Given the small number of studies which investigated the neural mechanisms of misophonia so far, these hypotheses remain tentative. Nevertheless, the increased functional connectivity of the lateral OFC allows linking misophonia to neural mechanisms of compulsive behaviour and behavioural inhibition for further investigations.

Additional insights into the role of the lateral OFC in misophonia are provided by studies on dysfunctions of the TAO network. Syndromes involving damage to this network - such as advanced Alzheimer and temporal lobe epilepsy - result in personality changes and other behavioural symptoms like aggressivity, disinhibition, deepened cognitive and emotional responses, antisocial traits (Catani *et al.*, 2013). Historically, the most famous case of traumatic damage to this network is that of Phineas Gage (Harlow, 1868; Thiebaut de Schotten *et al.*, 2015). Patients with traumatic brain injury resulting in OFC lesions show socially inappropriate behaviour, impulsivity, reduced control of the emotional response and perseveration (Thiebaut de Schotten *et al.*, 2015). Most elements of the symptomatology of TAO syndromes are reminiscent of the behavioural manifestations in misophonia: people with this condition are well aware of the disproportion between the stimuli and their compulsive emotional reaction, characterized by anger, irritation, stress, anxiety, which can yield to socially inappropriate behaviour and in extreme cases to aggressive responses (Rouw and Erfanian, 2018). In addition, a recent study examining a large sample of people reporting symptoms compatible with misophonia (N=575) showed that this condition has comorbid traits of obsessive-compulsive personality (26%) and mood disorders (10%) (Jager *et al.*, 2020). While our results highlight altered TAO network activity restricted to the lateral OFC cortex, these similarities between misophonia and neurological and neuropsychiatric syndromes grant further studies in the anatomical and functional connectivity between the regions connected within the TAO network.

### Direct vs. indirect auditory-limbic pathways underlying the emotional response in misophonia

Given that misophonia is associated with the exposure to sounds, it would be plausible to expect an alteration of the activity of auditory cortices during trigger sounds in people suffering from this condition. However, previous neuroimaging studies on misophonia failed to detect such effect in a whole-brain corrected analysis. Only one study examining a specifically selected region of the secondary auditory cortex (lateral superior temporal gyrus) reported significantly increased brain activity during trigger vs. generic aversive stimuli in people with misophonia (Schröder *et al.*, 2019).

Importantly, increased brain activity in the auditory cortex can be expected by variations in the physical properties of the sounds (such as volume) in neurotypical individuals, as well as in neurological patients with pathologically increased sensitivity to sound amplitude (Lanting *et al.*, 2008; Tyler *et al.*, 2014). However, misophonia appears to be unrelated to neurological syndromes or trauma, and audiological tests show that people with this condition have normal hearing capacities (Jager *et al.*, 2020). In addition, their emotional response appears to be independent from the volume of the trigger sounds.

In our analysis, we also failed to detect an increase in the brain activity of the auditory cortices of the superior temporal lobe. However the PPI analysis seeded in the MCC - whose activity was increased in trigger vs. generic aversive stimuli - revealed an increased task-dependent synchronization between MCC and the primary auditory cortex (A1) during the presentation of trigger stimuli.

The current model of misophonia proposes that this condition is associated with an altered interaction between the auditory and the limbic system (Jastreboff and Jastreboff, 2001). To our knowledge, the increased synchronization between MCC and A1 activity we report represents the first evidence of an atypical activity of primary auditory regions in misophonia. Importantly, such evidence is crucial to identify which among the possible direct and indirect neural pathways connecting auditory regions with the limbic system subserve the generation of the emotional response in misophonia. Given that people with misophonia most commonly qualify the emotional response as “extreme annoyance/irritation” or “anger/rage” (Rouw and Erfanian, 2018), and given the presence of monosynaptic connections between A1 and the amygdala (LeDoux *et al.*, 1991), such direct pathway could represent one suitable neural mechanism. Additionally, it was shown that both A1 and the amygdala show an increased activity and connectivity during generic aversive - such as nails scratching on a blackboard - vs. neutral sounds, and that their reciprocal directional connectivity encodes both physical properties and subjective valence (perceived unpleasantness) of these generic aversive sounds (Kumar *et al.*, 2012). A relation between physical properties (frequency range) of these sounds and their perceived unpleasantness of such sounds had been previously shown (Halpern *et al.*, 1986; Reuter and Oehler, 2011). However, when trigger sounds were contrasted with generic aversive sounds in people with misophonia, no significant increase in brain activity was detected in the amygdala (Schröder *et al.*, 2019), and only a marginally significant increased functional connectivity between the amygdala and the anterior insula - rather than the auditory cortex - was reported (Kumar *et al.*, 2017).

Together with this previous evidence, our result showing increased task-related functional connectivity between MCC, A1 and OFC during trigger stimuli presentation in people with misophonia make it unlikely that the neural pathways subserving the emotional and behavioural response in misophonia could be identified with a direct connection between the auditory cortex and the amygdala. Instead our results suggest that the emotional reaction to trigger sounds is subserved by indirect neural pathways connecting the auditory cortex with the dorsal anterior insula, the MCC and the OFC. Importantly, all these three regions are crucial in establishing high-order cognitive associations between sensory stimuli and emotional or behavioural responses. Specifically, the MCC is specialized for feedback-mediated decision making such as in reward/punishment anticipation and error selection (Vogt, 2019*a*), and together with the anterior insula is part of the saliency network which is crucial to integrate sensory, cognitive and interoceptive information to decide the most homeostatically relevant stimuli in determining behaviour (Seeley *et al.*, 2007).

### High-order vs. primary emotional associations in misophonia

One important open question in misophonia - related to the involvement of A1 in this behavioural condition - is whether the specificity of the stimuli triggering the excessive emotional response is due to the physical properties of the stimuli or to their subjective valence.

The absence of increased activity in A1 - processing the physical properties of auditory stimuli - and the involvement of OFC and MCC - crucial for the emotional estimation of the association between sensory stimuli and behavioural response - deposes against the possibility that the physical properties of the auditory stimuli alone can explain why these stimuli and not others elicit the disproportionate emotional response seen in misophonia.

The clinical literature provides large evidence to frame misophonia as a dysfunction related to high-order rather than basic sensory properties of trigger stimuli. This condition appears to be unrelated to neurological damage, and is not associated with hearing deficits (Schröder *et al.*, 2013). Instead its symptoms arise during childhood or adolescence, and increase with age (Rouw and Erfanian, 2018), which is compatible with the possibility that they rest on learned conceptual associations between specific innocuous sensory stimuli and emotionally distressing life experiences. Sufferers experience diminished or no distress when they produce trigger sounds themselves, and occasionally use this as a coping strategy (Edelstein *et al.*, 2013). The emotional reaction can be modulated by the social context, and amplified when trigger sounds are produced by significant others (Edelstein *et al.*, 2013; Bruxner, 2016). Symptoms can be alleviated by cognitive-behavioural therapy, while they are insensitive or even exacerbated by attempts to habituation (Bernstein *et al.*, 2013, Schröder *et al.*, 2017*a*; Aazh *et al.*, 2019).

Another important source of evidence for the involvement of high-order constructs in misophonia is represented by the comparison with hyperacusis: a condition characterized by hyperresponsivity to non-noxious auditory stimuli due to the amplified perception of their volume (Jastreboff and Jastreboff, 2001; Song *et al.*, 2014). While misophonia and hyperacusis present a certain degree of comorbidity and evoke similar negative emotional and behavioural reactions (Jastreboff and Jastreboff, 2015), hyperacusis is characterized by hyperaesthesia for generic sounds, modulated by their physical properties. Notably, hyperacusis appears to be associated with altered activity of the primary auditory cortex (Song *et al.*, 2014; Wong *et al.*, 2020). Instead misophonic triggers are very specific human-generated sounds, mostly produced with the mouth or the oropharyngeal tract and the intensity of the emotional response is independent from their volume.

The presence of an altered structural or functional circuitry linking the OFC with the insula, the amygdala and the MCC could represent the source of the behavioural manifestations of misophonia. Importantly, the fact that OFC is responsible for high-order, rather than primary associations between stimuli and emotional response, is compatible with the selectivity of misophonia for a limited set of stimuli, possibly related to previously learned negative associations (Peters and Büchel, 2010). Indeed increased task-related OFC connectivity - estimated by PPI analysis - has been previously recorded during tasks involving high-order associations between the stimulus and its emotional content, for instance with monetary reward (Noonan *et al.*, 2011), during self-motivation (Bengtsson *et al.*, 2009) and in the downregulation of pain associated with monetary gain (Becker *et al.*, 2017), but not when exposed to intrinsically emotionally loaded stimuli such as pain (Perini *et al.*, 2020). Interestingly, it was shown how the OFC functional connectivity (with the amygdala) is instead related to the reappraisal of stimuli containing negative basic emotions (Banks *et al.*, 2007), which requires establishing high-order associations to downregulate the primary emotional effect of the stimuli.

In summary, all the available evidence - including our results - lead to hypothesize that selectivity of misophonia for specific sounds, and the emotional reaction triggered by them, cannot be explained only considering the physical properties of the stimuli, but rather by high-order cognitive associations which are potentially related to previously established negative associations with the same stimuli. On the other hand, there is strong evidence that auditory stimuli commonly serve as 'triggers' in misophonia, rather than stimuli in other modalities. This specific relationship between the auditory system and misophonia still is largely unexplained. For example, an accurate analysis of the specific frequencies of the most common trigger stimuli - similar to what that carried out for generic stimuli by (Kumar *et al.*, 2012), has not yet been performed, and it will be crucial to understand whether a specific mixture of frequencies plays a role in understanding the specificity of trigger vs. generic aversive sounds.

### Differences between seed-based correlation and psychophysiological interaction (PPI)

One important previous neuroimaging result in misophonia showed an increased task-dependent functional connectivity between the anterior dorsal insula and the medial regions of the default mode network (DMN), which is involved in internally directed high-order information processing such as social and autobiographical memory (Callard and Margulies, 2014; Margulies *et al.*, 2016). It was suggested that this increased functional coupling could signal a difficulty in people with misophonia to dissociate the sensory stimulation of the trigger sounds from its contextual associations and memories (Kumar *et al.*, 2017). In our analysis we used the same region of interest - the anterior right insula from the SPM12 neuromorphometrics toolbox - to carry out a PPI analysis, however we couldn’t replicate the previously reported increased insula/DMN functional connectivity.

While this could be due to differences in the paradigm (see above “Comparison with previous neuroimaging studies”), it is important to stress that the two methods used here and in (Kumar *et al.*, 2017) to probe task-dependent changes in functional connectivity isolate different properties of the interaction between brain regions. PPI estimates the effects of the interaction between the task and the seed region *over and above* the correlation between the activity of seed and target region (Friston *et al.*, 1997; O’Reilly *et al.*, 2012; Di *et al.*, 2020) - e.g. anterior insula and DMN. Instead the latter correlation is precisely what is reported in the weighted seed-based connectivity (wSBC) used in (Kumar *et al.*, 2017). While in wSBC only the time points where the stimulus is presented are considered - and in this sense the retrieved correlation is task-based - it cannot be excluded that the results are due not only to the task, but also to other sources of fMRI signal, such as resting-state or the effect of a third region driving the correlation between seed and target. While PPI is also affected by the latter issue, in PPI explicitly modelling out the signal of the seed region ensures that the change in functional connectivity is a direct function of task-related activity, and cannot be attributed to resting-state signal or task-unrelated changes in signal-to-noise ratio (Friston, 2011; O’Reilly *et al.*, 2012).

This difference between the interpretation of wSBC and PPI results does not in any way compromise or diminish the importance of the result by Kumar and colleagues. Instead, it opens the possibility that the observed task-dependent increased connectivity between the anterior insula and the DMN might be due to *structural* differences in the functional organization of resting-state networks in people with misophonia; an hypothesis which is also supported by the co-localization of this functional atypicality with a higher myelination of the anterior cingulate cortex (Kumar *et al.*, 2017). A crucial test for this hypothesis will be to compare the spontaneous brain activity in the DMN in people with and without misophonia: an analysis for which specific fMRI data not available in our dataset needs to be acquired.

Related to this, also our results suggest that there might be differences in spontaneous brain activity in misophonia, as the OFC did not display a change in the overall brain activity due to trigger vs. aversive stimuli, but only in its functional connectivity with other brain regions where trigger stimuli prompted changes in overall brain activity (SMA, MCC and vlPMC). As mentioned above in the Methods, the results of PPI can refer either to (1) the task-dependent modulation of the interaction between region A and region B, or to (2) the effect of the modulatory activity of region A onto region B during a task (See Fig. 5 in (Friston *et al.*, 1997)). In our case the lack of increased activity in the OFC during trigger vs. stimuli suggest that situation (2) is more likely to reflect the brain mechanisms influencing the behavioural response in misophonia, and that in this condition OFC exerts a *tonic* influence on SMA, MCC and vlPMC which results in higher activity during trigger stimuli in the latter regions. Again, an analysis of resting-state fMRI activity in misophonia would represent an important step to disentangle between the two possible situations.

## Conclusions

In the present fMRI study we investigated the neural mechanisms underlying the disproportionate negative emotional response elicited by trigger stimuli in misophonia. Besides replicating previous evidence of the increased insular, premotor and mid-cingulate activity in misophonia during trigger stimuli, we found that in people with this condition trigger stimuli prompt an increased synchronization of premotor and mid-cingulate activity with the lateral OFC.

The involvement of the OFC, crucial for establishing and revising specific experience-dependent cognitive-emotional associations, represents the first evidence that allows to shed light on the selectivity of misophonia for particular innocuous auditory stimuli. Specifically, the increased functional connectivity of this region with premotor and mid-cingulate cortex might reflect a dysfunction in the neural mechanisms required to reappraise a previously learned negative emotional association with specific sounds.

The increased synchronization between these regions could furthermore promote the establishment and reinforcement of a functional unit whose activity is automatically triggered by an appropriate stimulus, generating a compulsive - and therefore difficult to prevent - emotional and behavioural response. While so far, the critical role of the OFC in creating persistent responses at a particular task or in a particular situation has so far been shown in other behavioural conditions (addiction, compulsive behavior, (impaired) decision making) such framework can help explain the persistent emotional and behavioral responses that are so characteristic of misophonia.

We also report the first evidence relating misophonia with the primary auditory cortex. This region did not show an increased activity to trigger vs. aversive sounds, but rather an increased synchronization with the mid-cingulate during trigger sounds. This finding, together with the increased interaction between mid-cingulate and OFC cortex, supports the hypothesis that the emotional reaction in misophonia is unlikely to reflect a direct response of the limbic system to the physical properties of the sounds alone. Rather it suggests that the emotional response is mediated by a complex network of brain regions processing the association between specific sounds and their subjective cognitive, emotional and autonomic valence in people with misophonia.

To further clarify the neural mechanisms of misophonia, we propose that further studies should focus on the direct comparison between misophonia and hyperacusis, since the latter condition presents the most similar pattern of symptoms, with the exception of the stimulus selectivity in misophonia. For these reasons, a sample of people with hyperacusis might represent the best control group to precisely determine the brain mechanisms which characterize misophonia.

