## Supplementary Materials for "Increased orbitofrontal connectivity in misophonia"

| | |
|---|---|
| 1 | I feel angry when I see and / or hear a trigger<br>*Ik voel me boos als ik een trigger hoor en/of zie* |
| 2 | I am disgusted when I hear and / or see a trigger<br>*Ik walg als ik een trigger hoor en/of zie* |
| 3 | Sometimes I behaved aggressively because of my misophonia<br>*Ik heb me weleens agressief gedragen door mijn misofonie* |
| 4 | I get stressed or panic when I hear and / or see a trigger<br>*Ik word gestrest of paniekerig als ik een trigger hoor en/of zie* |
| 5 | If I hear and / or see a trigger, I would like to hurt the person who causes it<br>*Als ik een trigger hoor en/of zie dan zou ik de persoon die het veroorzaakt iets willen aandoen* |
| 6 | I feel my muscles tense when I hear and / or see a trigger<br>*Ik voel mijn spieren spannen als ik een trigger hoor en/of zie* |
| 7 | I feel physically uncomfortable when I hear and / or see trigger<br>*Ik voel me fysiek onaangenaam als ik een triggers hoor en/of zie* |
| 8 | I feel panic / anxiety / stress at the idea that a trigger may occur<br>*Ik voel paniek/angst/stress bij het idee dat een trigger zich kan voordoen* |
| 9 | If my trigger is present, I can only think about that<br>*Als mijn trigger aanwezig is, kan ik alleen nog maar daar aan denken* |
| 10 | My misophonia limits my functioning in daily activities (work / relationship / daily life)<br>*Mijn misofonie beperkt mij in mijn functioneren (werk/relatie/dagelijks leven)* |
| 11 | The triggers lead me to avoid social situations<br>*De triggers leiden ertoe dat ik sociale situaties vermijd* |
| 12 | I have suicidal thoughts because of my misophonia<br>*Ik heb suïcidale gedachten vanwege mijn misofonie* |
| 13 | I have sometimes hurt myself in order to deal with my misophonia<br>*Ik heb mijzelf weleens pijn gedaan, om met mijn misofonie om te kunnen gaan* |
| 14 | I am afraid to lose control of myself because of misophonia<br>*Ik ben bang door mijn misofonie de controle over mijzelf te verliezen.* |
| 15 | Misophonia has a strong effect on my life<br>*Misofonie heeft een groot effect op mijn leven* |

Misophonia questionnaire used to select the participants for the present experiment, among the 232 people we contacted, who referred symptoms potentially related with misophonia. Besides the first question ("I know what misophonia is"), people were requested to report their agreement/disagreement with each statement on a 5-points scale ranging from 1 ("Strongly disagree") to 5 ("Strongly agree"). We considered for the neuroimaging experiment only people reporting misophonic symptoms whose response at each question was equal or greater than 3. We used the same questionnaire to check for absence of strong indication of misophonia (any response equal or smaller than 3). The questionnaire was created by aggregating the commonalities among a number of previous questionnaires for misophonia symptoms proposed in the literature or in the clinical practice (Fitzmaurice 2010; Edelstein et al. 2013; Schröder et al. 2013; Bauman 2015; Dozier 2015; Wu et al. 2014).

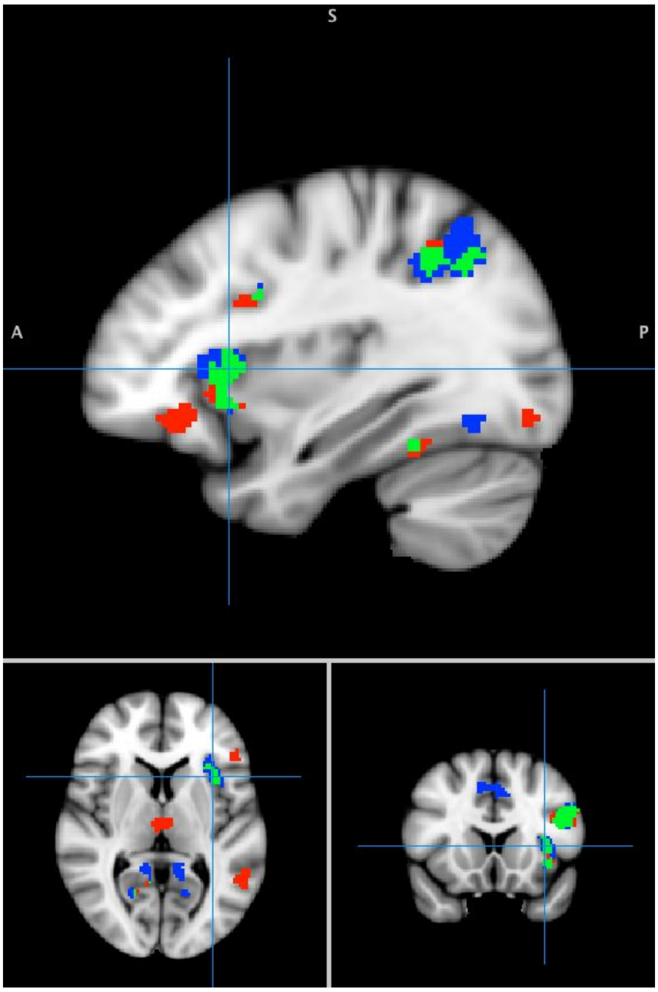


**Supplementary Figure 1 - Regions showing higher intensity in either groups during the description of the incoming audio-visual, with respect to the audio-visual itself**. The overlay shows all voxels above a threshold of $Z > 3.1$ and corrected for multiple comparison at a cluster significance (using Gaussian Random Fields) of $p=0.05$ (Worsley 2001). Red indicates the result in people with misophonia, blue indicates the result in control participants, while green indicates the voxels where the effect was significant in either group. The comparison of these parameter estimates between groups did not lead to significant effects in any brain voxel.
